# Local organization of spatial and shape information in the primate prefrontal cortex

**DOI:** 10.1101/2023.08.26.554962

**Authors:** Yunyi Sun, Wenhao Dang, Rye G. Jaffe, Christos Constantinidis

**Affiliations:** Department of Biostatistics, Vanderbilt University Medical Center, Nashville TN 37203, USA; Department of Biomedical Engineering, Vanderbilt University, Nashville TN 37235, USA; Neuroscience Program, Vanderbilt University, Nashville TN 37235, USA; Department of Ophthalmology and Visual Sciences, Vanderbilt University Medical Center, Nashville TN 37232, USA

## Abstract

The current understanding of sensory and motor cortical areas has been defined by the existence of topographical maps across the brain surface, however, higher cortical areas, such as the prefrontal cortex, seem to lack an equivalent organization, and only limited evidence of functional clustering of neurons with similar stimulus properties is evident in them. We thus sought to examine whether neurons that represent similar spatial and object information are clustered in the monkey prefrontal cortex and whether such an organization only emerges as a result of training. To this end, we analyzed neurophysiological recordings from male macaque monkeys before and after training in spatial and shape working memory tasks. Neurons with similar spatial or shape selectivity were more likely than chance to be encountered at short distances from each other. Some aspects of organization were present even in naïve animals, however other changes appeared after cognitive training. Our results reveal that prefrontal microstructure automatically supports orderly representations of spatial and object information.

## INTRODUCTION

The prefrontal cortex (PFC) has long been recognized as the seat of higher cognitive functions, based on a plethora of evidence, including lesion, anatomical, and neurophysiological experiments in non-human primates (Goldman-Rakic 1988). A population of prefrontal neurons generate persistent activity selective for the properties of stimuli held in memory (Fuster and Alexander 1971; Funahashi et al. 1989; Miller et al. 1996; Mendoza-Halliday et al. 2014). Computational models typically simulate such persistent activity by networks of units with recurrent connections; units with similar tuning for stimulus properties are more strongly connected with each other and this organizations allows activity initiated by a specifical stimulus to be maintained in the network during working memory (Wimmer et al. 2014; Hart and Huk 2020).

The prefrontal cortex lacks an obvious retinotopic organization (Kiani et al. 2015; Leavitt et al. 2017), or other type of systematic topography of sensory information that would be analogous to the orderly arrangement of units in these artificial networks – or in the sensory and motor cortices. Only some local organization has been previously described in the prefrontal cortex, for spatial information, with nearby neurons representing the same or adjacent spatial locations (Constantinidis et al. 2001; Arnsten 2013). It has been suggested that much of the information represented by prefrontal neurons is the result of training, reflecting task demands. For example, single neurons are thought to represent jointly object and spatial information, only if trained in a task that requires such conjunctive representations (Rao et al. 1997). It is unclear therefore how neurons with similar selectivity can organize into functional circuits.

Newer theories of working memory, motivated primarily by results of human imaging studies have also proposed that working memory information is represented in the sensory cortex, instead (Harrison and Tong 2009; Albers et al. 2013; Ester et al. 2013; Xing et al. 2013; Sreenivasan et al. 2014). According to this viewpoint, prefrontal cortex plays a control or supervisory role, perhaps highlighting the locations of remembered stimuli, rather than maintaining their identity in working memory itself (Curtis and D’Esposito 2003; Tong and Pratte 2012; D’Esposito and Postle 2015). It is possible, therefore, that any local organization within the prefrontal cortex is limited to spatial information, alone.

We were thus motivated to examine the local organization of prefrontal neurons for the maintenance of spatial and object information in memory. We first replicated the analysis of spatial information of prior studies (Constantinidis *et al*. 2001), from a different set of experimental data. We then applied the same analysis to object information. Additionally, we examined how this information is altered by training to perform a working memory task, by analyzing data recorded from monkeys naïve to training, and after they were trained to perform spatial and object working memory tasks.

## METHODS

### Animals

Data obtained from six male rhesus monkeys (*Macaca mulatta*), ages 5–9 years old, weighing 5–12 kg, as previously documented (Riley et al. 2018), were analyzed in this study. None of these animals had any prior experimentation experience at the onset of our study. Monkeys were either single-housed or pair-housed in communal rooms with sensory interactions with other monkeys. All experimental procedures followed guidelines set by the U.S. Public Health Service Policy on Humane Care and Use of Laboratory Animals and the National Research Council’s Guide for the Care and Use of Laboratory Animals and were reviewed and approved by the Wake Forest University Institutional Animal Care and Use Committee.

### Experimental setup

Monkeys sat with their heads fixed in a primate chair while viewing a monitor positioned 68 cm away from their eyes with dim ambient illumination and were required to fixate on a 0.2° white square appearing in the center of the screen. In order to receive a liquid reward (typically fruit juice), the animals maintained fixation on the square while visual stimuli were presented either at a peripheral location or over the fovea. Any break of fixation immediately terminated the trial and no reward was given. Eye position was sampled at 240 Hz, monitored on line, and recorded with an infrared eye position scanning system (model RK-716; ISCAN, Burlington, MA). The visual stimulus display, monitoring of eye position, and synchronization of stimuli with neurophysiological data was performed with in-house software (Meyer and Constantinidis 2005) implemented on the MATLAB environment (Mathworks, Natick, MA), utilizing the Psychophysics Toolbox (Brainard 1997).

### Pre-training task

Following a brief period of fixation training and acclimation to the stimuli, monkeys were required to fixate on a center position while stimuli were displayed on the screen. The stimuli shown in the pre-training passive spatial task were white 2° squares, presented in one of nine possible locations arranged in a 3 × 3 grid with 10° distance between adjacent stimuli. The stimuli shown in the pre-training passive feature task were white 2° geometric shapes drawn from a set comprising a circle, diamond, the letter H, the hashtag symbol, the plus sign, a square, a triangle, and an inverted Y-letter. These stimuli were also presented in one of nine possible locations arranged in a 3 × 3 grid with 10° distance between adjacent stimuli.

Presentation began with a fixation interval of 1 s where only the fixation point was displayed, followed by 500 ms of stimulus presentation (referred to hereafter as cue), followed by a 1.5 s “delay” interval (referred to hereafter as delay1) where, again, only the fixation point was displayed. A second stimulus (referred to hereafter as sample) was subsequently shown for 500 ms. For the spatial stimulus set, this second stimulus would be either identical or diametrically opposite in location to the initial stimulus. For the feature set, this second stimulus would appear in the same location to the initial stimulus and would either be an identical shape or the corresponding non-match shape (each shape was paired with one non-match shape). Only one nonmatch stimulus was paired with each cue, so that the number of match and nonmatch trials would be balanced in each set. In both the spatial and feature task, this second stimulus display was followed by another “delay” period (delay2) of 1.5 s where only the fixation point was displayed. Successfully maintaining fixation until the end of the second delay period resulted in reward. The location and identity of stimuli was of no behavioral relevance to the monkeys during the “pre-training” phase, as fixation was the only necessary action for obtaining reward.

### Post-training task

Four of the six monkeys were trained to complete spatial and feature working memory tasks. These tasks involved presentation of identical stimuli as the spatial and feature tasks during the “pre-training” phase, but now monkeys were required to remember the spatial location and/or shape of the first presented stimulus, and report whether the second stimulus was identical to the first or not, via saccading to one of two target stimuli (green for matching stimuli, blue for non-matching). Each target stimulus could appear at one of two locations orthogonal to the cue/sample stimuli, pseudo-randomized in each trial.

### Surgery and neurophysiology

The animals were initially implanted with a headpost device under inhalant anesthesia. Opioid analgesics were administered after the surgery and the animals were allowed to recover for at least three weeks before behavioral sessions began. They were then exposed to the visual stimuli, as described in the pre-training phase. A second surgery was subsequently performed, in which a 20 mm diameter craniotomy over the PFC was performed and a recording cylinder was implanted over the site. The location of the cylinder was visualized through anatomical magnetic resonance imaging (MRI) and stereotaxic coordinates post-surgery. Electrode penetrations were aligned to the MRI image and mapped onto the cortical surface.

### Neuronal recordings

Neural recordings were carried out both before and after training in the working memory tasks. The recordings analyzed here were previously used to determine the properties of neurons before and after training in different PFC subdivisions and additional details can be found in previous reports (Meyer et al. 2011; Riley *et al*. 2018). Extracellular recordings were performed with multiple epoxylite-coated tungsten microelectrodes, with a 100 μm diameter and 1–4 MΩ impedance at 1 kHz (FHC, Bowdoin, ME). A Microdrive system (EPS drive, Alpha-Omega Engineering) advanced arrays of up to 8-microelectrodes, spaced 0.2–1.5 mm apart, through the dura and into the PFC. The signal from each electrode was amplified and band-pass filtered between 500 Hz and 8 kHz while being recorded with a modular data acquisition system (APM system, FHC, Bowdoin, ME). Waveforms that exceeded a user-defined threshold were sampled at 25 μs resolution, digitized, and stored for off-line analysis. Neurons were sampled in an unbiased fashion, collecting data from all units isolated from our electrodes, with no regard to the response properties of the isolated neurons. A semi-automated cluster analysis relied on the KlustaKwik algorithm, which applied principal component analysis of the waveforms to sort recorded spike waveforms into separate units (Harris et al. 2000). To ensure a stable firing rate in the analyzed recordings, we identified recordings in which a significant effect of trial sequence was evident at the baseline firing rate (ANOVA, p < 0.05), e.g., due to a neuron disappearing or appearing during a run, as we were collecting data from multiple electrodes. Data from these sessions were truncated so that analysis was only performed on a range of trials with stable firing rate. Less than 10% of neurons were corrected in this way. Identical data collection procedures, recording equipment, and spike sorting algorithms were used before and after training in order to prevent any confounds.

### Data analysis

Analysis was implemented with the MATLAB computational environment (R2020-2023, Mathworks, Natick, MA). To obtain tuning similarity analysis by optimal stimuli, we first identified neurons with significant tuning for the stimulus set being examined. A one-way ANOVA at the α=0.05 significance level was employed, separately for each epoch of the task (fixation, cue, delay1, sample, delay2 in Fig. 1). The preferred stimulus in each task epoch was then defined for the neuron as the stimulus (location/shape) that elicited the highest average firing rate in the corresponding epoch. We examined pairs of nearby neurons recorded simultaneously from separate microelectrodes, placed together at distances of 0.2 – 1.5 mm from each other. For each task epoch, we identified such pairs both neurons of which were selective for the stimuli, and determined if they had the same preferred (best) stimulus or not. The proportions of such neuron pairs among all tuned neuron pairs were determined in each epoch and we used a nonparametric test to test the statistical significance of this proportion. More specifically, we randomly paired cells in our pool that were not recorded simultaneously (from nearby locations), and then repeated the same analysis described above to determine a simulated proportion of cells with the same preference. This random pairing procedure was repeated 1000 times to get a full null distribution. We then counted the number of occurrences in the null distribution for more extreme values compared to the empirical value and divided that number by 1000 to get a probability measure, which would be the p-value of the empirical measurement. For comparing two proportions in the pre- and post-training conditions, we used calculated a z-score as follows:

**Figure 1.**
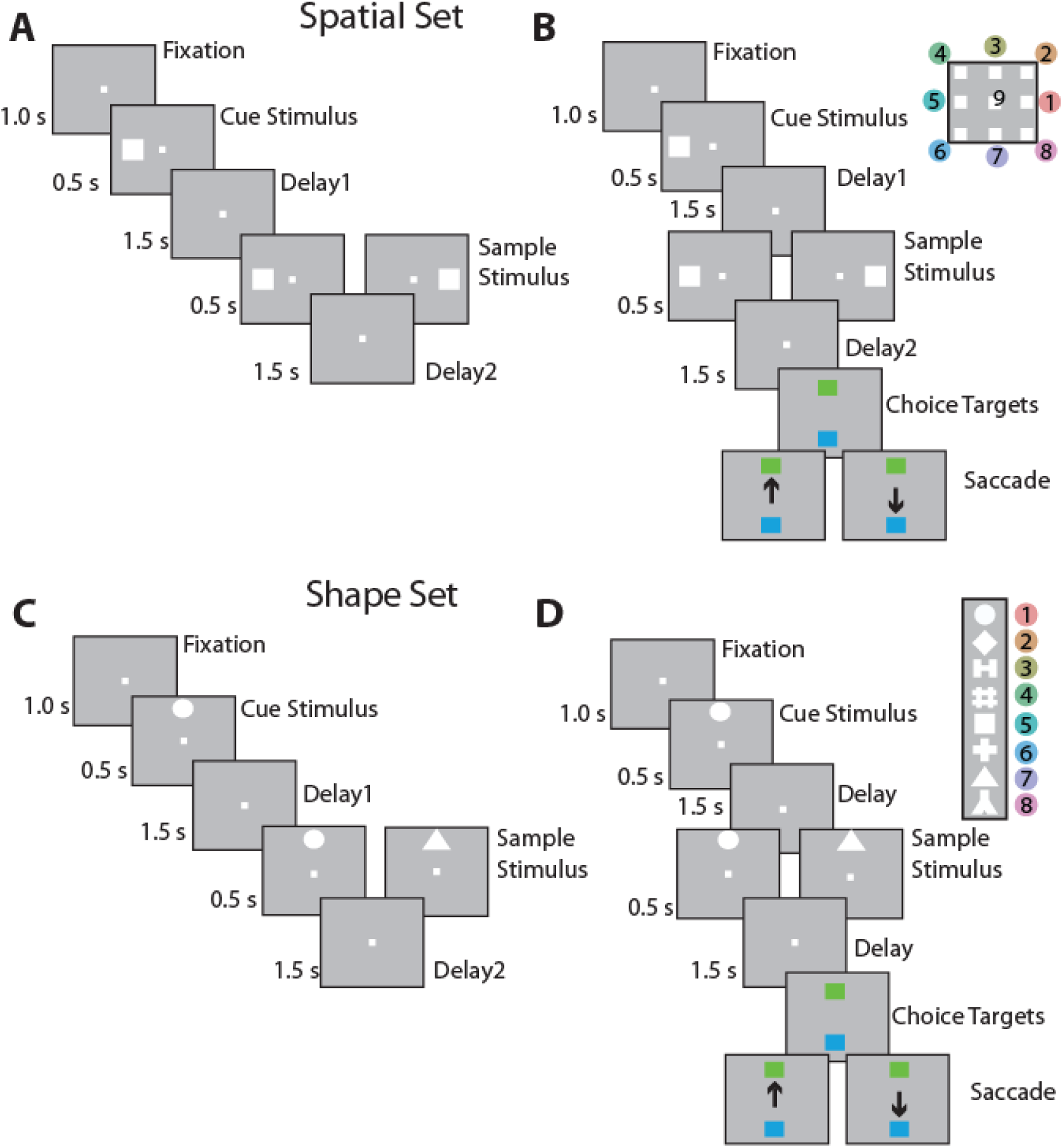
Task structure and stimuli used. A. Passive, spatial task. B. Active spatial task. nine possible cue locations in a session shown in the inset. The peripheral locations are numbered from 1-8, for reference in other figures. The animals were required to maintain center fixation throughout both active and passive task trials. At the end of active tasks trials however, monkeys were required to make a saccade to a green target if the stimuli matched or to a blue target if the stimuli did not match. C-D. Passive and active shape task. Eight possible shapes in a session are shown in the inset, numbered from 1-8, for reference in other figures. Stimuli in both tasks extended 2 degrees of visual angle. Modified based on (Riley *et al*. 2017).

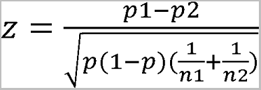, where 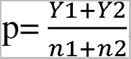, n1 and n2 are the sample sizes of the two populations, p1 and p2 are the proportions of significant cells, and Y1 and Y2 are counts of significant cells. The null hypothesis was rejected for values of z≥1.96.

In addition to determining the best stimulus of each pair, we also analyzed the tuning similarity of nearby neurons by quantifying the Euclidean distance defined by their average response to all stimuli used in each task. More specifically, for a given neuron, a trial-average firing rate vector was created by averaging the trials recorded with each stimulus, resulting in a 9 by 1 vector for the spatial test, and an 8 by 1 vector for the feature set. Each neuron’s firing rate vector for a certain task epoch was then normalized by the highest value in the vector, to control for firing rate difference between neurons. For a given nearby neuron pair, and for a given task epoch, we calculated the Euclidean distance in the 9-(or 8-) dimensional space using the built-in MATLAB function “pdist2”. If both cells in the pair displayed selectivity in more than one task epoch, the averaged tuning distance for all task epochs in which significant selectivity was present was used. We used a nonparametric test for testing the statistical significance of the Euclidean distance measurement, by comparing the empirical measurement to the results from two control conditions. We again randomly paired selective cells or non-selective neurons from different recordings, without having been recorded simultaneously. For a given neuron in this cell pool, we randomly chose one of the four epochs to get data from. This is the first source of our randomness introduced to the bootstrap analysis. Then we constructed a random cell pair pool containing all possible combinations of cells in the pool. And lastly we randomly drew the same number of pairs of cells as in the empirical pair dataset to calculate one value of the mean Euclidean distance for the null distribution. We had 1000 such iterations to generate the full null distribution. By comparing real neighboring selective cells to random paired selective cells, we could detect whether a physical clustering effect was present, and by comparing to the empirical result from non-selective pairing group we could confirm that the difference in Euclidean distance we found was not due to the difference in the statistics between selective and nonselective cells.

To visualize the representational similarity of different stimuli at the population level, we used the MATLAB function “linkage” and “dendrogram”. Specifically, we created an nine by N matrix (or eight by N matrix for the shape stimuli), where N is the number of unique neurons in the empirical neighboring-neuron pool as described above. Each element in the matrix represents the average neural response to 9 (or 8) stimuli. We then use the MATLAB function “pdist2” to create an 9 x 9 (or 8 by 8) pairwise distance matrix representing the pairwise distance between the neuronal representation of the stimuli. Lastly, the distance matrix served as the input to the MATLAB “linkage” function to create a hierarchical cluster tree. The representational similarity measurement was compared to two predictions based on perceptual similarity and pairing relationships of stimuli, for both the spatial and the feature tasks. Specifically, for the spatial task, the perceptual similarity is determined by the Euclidean distance between the locations. The further apart two locations are, the less similar they are perceptually. As for the pairing relationship prediction, we assume that the stimuli used as match/nonmatch pairs should be further apart in the neural representational space as well. For the spatial task, pairing based similarity is identical to perceptual based similarity, since locations symmetric to the center were paired as match-nonmatch pairs in the task. For the feature task, perceptual similarity was determined by using a computational model, ShapeComp (Morgenstern et al. 2021) that predicts human shape similarity judgements. The model projects 8 shapes used in task into a 2D perceptual similarity space, and the Euclidean distance in this space between shapes was used as the perceptual distance between shapes. We then constructed the dissimilarity matrix between locations/shapes and calculated the distance between the two dissimilarity matrices as 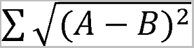, where A and B represent range-normalized dissimilarity matrices. To statistically test if the empirical pattern is more similar than chance to either prediction, a permutation test was performed. In each iteration, the location/shape labels were shuffled, followed by the same dissimilarity matrix distance measurement procedure as the empirical data. This procedure was repeated for 1000 times to construct a null distribution. The p value of the empirical measurements was determined by the number of occurrences of more extreme values in the null distribution.

## RESULTS

### Database

Extracellular neurophysiological recordings were collected from the lateral PFC of six male monkeys before and after they were trained to perform the match/nonmatch working memory tasks (Meyer *et al*. 2011; Riley *et al*. 2018). The task required them to view two stimuli appearing in sequence with an intervening delay period between them, and after a second delay period to report whether or not the second stimulus was identical to the first (Fig. 1). The two stimuli could differ in terms of their location (spatial task, Fig. 1A-B) or shape (feature task, Fig. 1C-D). If the second stimulus was identical to the first, monkeys were required to saccade towards a green choice target that appeared at the end of the trial. Otherwise, they had to saccade to a blue target. The locations of the choice targets varied randomly from trial to trial and the monkey could not plan a motor response prior to their appearance.

We collected neurophysiological data before and after the monkeys learned to perform these tasks. A total of 1617 cells from six monkeys were recorded while the animals viewed the spatial stimuli passively prior to training, and 1493 cells from five monkeys were recorded while the animals viewed the feature stimuli passively, prior to training to perform the task (“pre-training” phase). A total of 1104 cells from three of the same monkeys were collected after the monkeys were trained and were performing the spatial working memory task. Another 1091 cells from two of the same monkeys were collected after the monkeys were trained and performed the feature working memory tasks (“post-training” phase). Anatomical penetrations in each subject are shown in Supplementary Figure S1.

### Organization of Spatial Selectivity

We first identified neurons tuned for the spatial location of stimuli, evidenced by significant selectivity for the nine spatial stimuli (1-way ANOVA, p<0.05). Percentages of neurons that exhibited such significant selectivity at different time epochs of the task (plus the baseline, fixation period, as a control) are shown in Fig. 2. Spatial selectivity during the first delay period greatly increased after training. We observed both a higher proportion of selective cells (Fig. 2A-B, z-score based proportion test, p=4.39×10^-5^) and an overall higher degree of selectivity (Fig. 2C, two sample t-test, p=1.09×10^-4^), in agreement with results reported previously (Riley *et al*. 2018)

**Figure 2.**
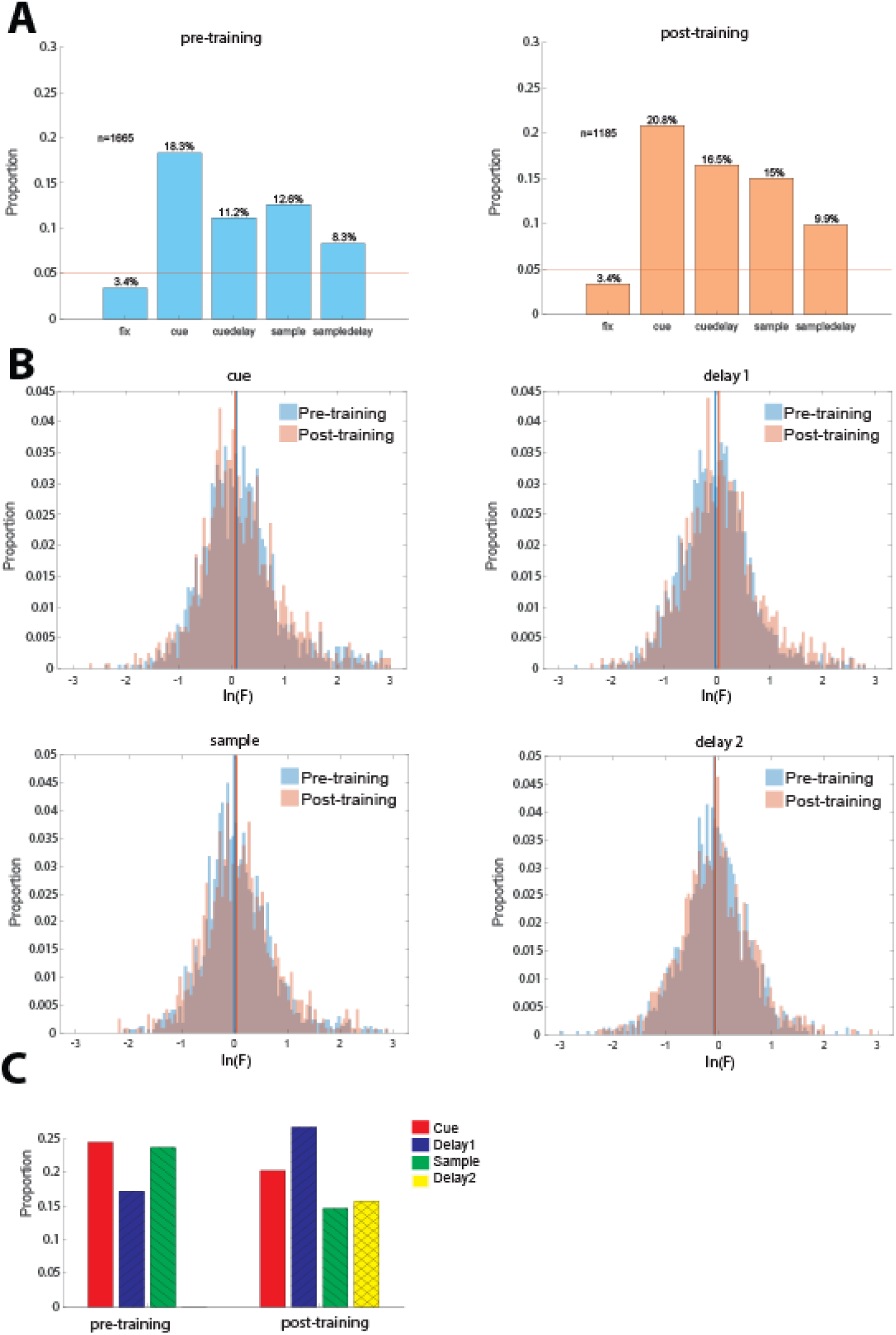
A. Proportion of neurons exhibiting significant spatial selectivity at different task epochs. Left. Selective neurons prior to training. Sample size n=1661. Right. Selective neurons post-training (n=1185 neurons). Training significantly increased the proportion of selective cells in the first delay epoch (Pre vs. Post z-score proportion test, Cue p=0.093, Delay1 p=4.39×10^-5^, Sample p=0.065, Delay2 p=0.161). B. Distributions of natural logarithm of F-values in the ANOVA test for location selectivity. Colored vertical lines represent median of distributions for corresponding training status.. C. Proportion of neurons with the same preferred cue location at different task intervals.

We relied on this dataset to examine how neurons selective for spatial stimuli are arranged in local prefrontal circuits. We thus examined 8139 pairs of neurons recorded simultaneously from different microelectrodes, spaced 0.2 – 1.5 mm apart from each (mean distance = 0.77 mm). For 6.13% of all examined pairs, (499 pairs total, 222 recorded before and 277 recorded after training), both neurons exhibited significant selectivity in the same task epoch. We could thus test whether neurons that had the same spatial preference (best response for the same location) were encountered more often than would be expected by chance (Fig. 2C), which was evaluated using a bootstrap task, randomly pairing neurons recorded at different sessions and random positions. We indeed determined that neurons with the same preference were significantly higher than expected at multiple task epochs: cue, and sample period, prior to training (p<0.001 in each case), and cue and delay period after training (p=0.012 and p<0.001, respectively).

This analysis considers only the best location, however, the entire tuning function of a neuron provides a more complete picture of neuronal tuning. We thus refined our analysis to consider the vector defined by firing rates for all eight locations for each neuron. We then computed the Euclidean distance between the vectors of two neurons, examining again only task epochs in which both neurons exhibited significant spatial selectivity. For pairs with significant selectivity in more than one task epoch, we computed distances in each epoch and then averaged them to obtain a single distance value for each neuron. To ensure that neurons with higher firing rates did not appear to exhibit longer distances, we normalized firing rates before computing distances. The mean distance computed in this manner was 1.02 prior to training and 1.11 after training (Fig. 3A). Both values were significantly lower than would be expected by chance (permutation test, p<0.001 compared to both control groups in each case).

**Figure 3.**
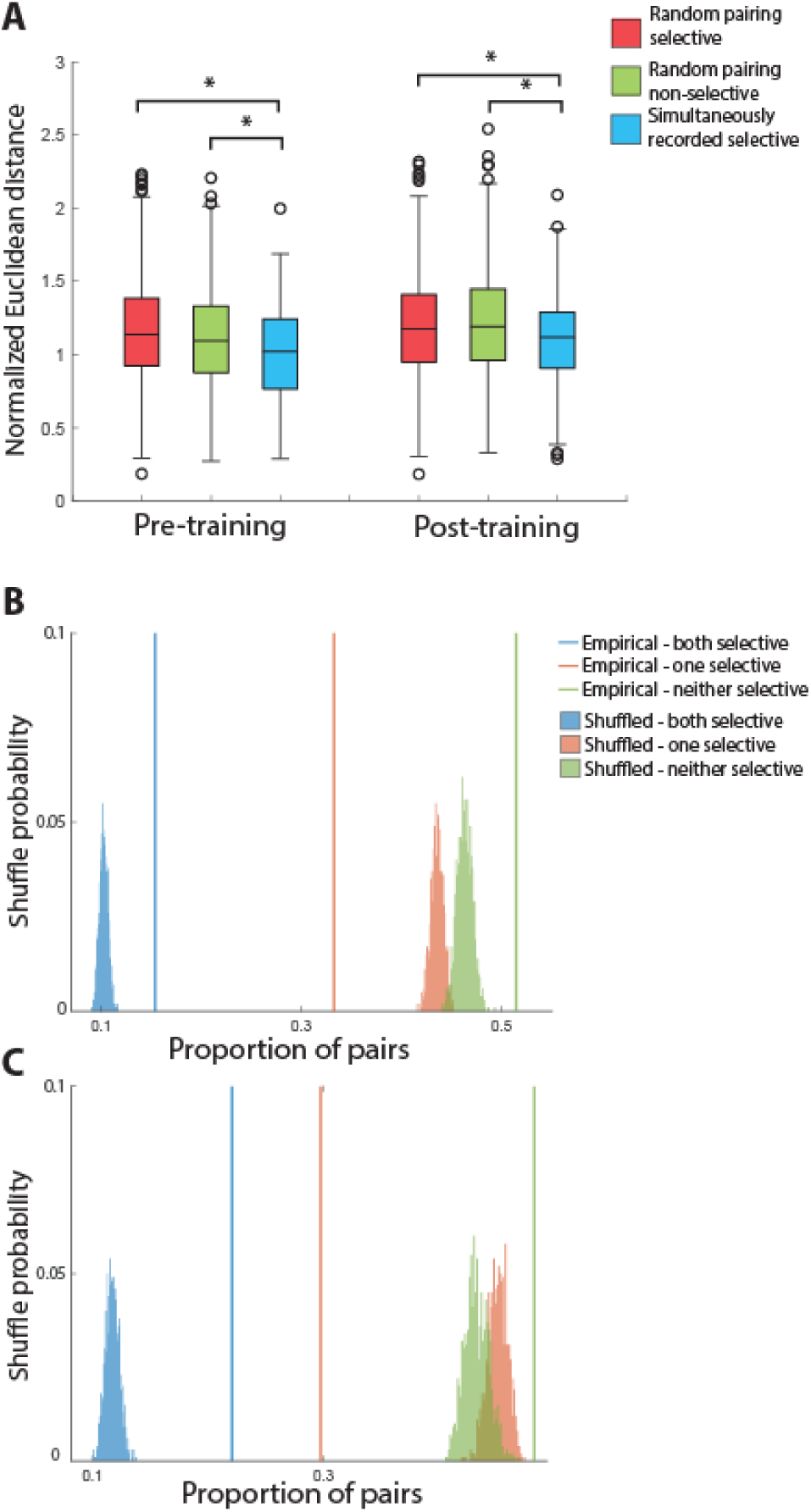
A. Normalized Euclidean distance of the representations of spatial stimuli among pairs of neighboring prefrontal neurons. The box borders represent the interquartile range (IQR) of the null distribution generated randomly. Whisker length indicates 1.5 interquartile range (IQR). Open circles are outliers in the null distribution. White boxes represent distributions from simultaneously recorded neighboring selective cell pairs in the same session. Black and gray boxes represent two control conditions, where either selective cells or non-selective cells were randomly paired, despite where two cells were from the same session. The representational distance between nearby neurons is smaller than the expected chance level. B. Permutation test for clustering selective cells in the pre-training spatial task. Colored vertical lines indicated the proportion of selective/selective, selective/nonselective, nonselective/nonselective pairs drawn from nearby (distance <1.5mm), simultaneously recorded neurons. This empirical proportion was compared to a null distribution constructed by shuffling session labels. C. Selective clustering permutation test for post-training data. Plotting conventions are the same as panel B.

This analysis demonstrated that nearby neurons were more likely to represent similar locations. We also determined the distance between neurons recorded at any distance of each other, based on the anatomical coordinates of each penetration. This analysis still revealed an overall positive relationship (Supplementary Figure S2).

To better characterize the local organization of local circuit selectivity, we further asked whether spatial location selective neurons are clustered in PFC, regardless of whether they prefer the same location. We calculated the proportion of selective/selective, selective/nonselective, and nonselective/nonselective cell pairs for all of our simultaneously recorded, neighboring cells (distance <1.5mm). We then compared the empirical results to a null distribution constructed by shuffling the session labels of the cells, to remove the relationship of neighboring locations in the cortex. This analysis revealed that simultaneously recorded, nearby cell pairs have much higher proportion of selective/selective pairs compared to the label-shuffled result, and this was true both before (Fig. 3B) and after training (Fig. 3C). The result suggests that spatially selective cells are clustered in the cortex.

The pattern of responses that nearby neurons exhibited with respect to different stimuli also allowed us to determine how different spatial locations are represented across different neurons. We thus performed a representational similarity analysis to visualize topography of stimulus representation on a population level, as well as a hierarchical clustering analysis, grouping stimuli depending on how similar responses to them were across neurons. Specifically, we first asked if there exist a ‘default’ representation of stimuli in PFC, so that the topography of stimuli representation reflects the perceptual similarity of stimuli; and second, if training will change the topography of stimuli representation. We first constructed a representational dissimilarity matrix based on location proximity and match/nonmatch pairing relationship of stimuli (Fig. 4A). The matrix distance was calculated between the real neural responses and this hypothetical prediction (Fig. 4B,C). We found that the population representation during the cue period already resembled the pattern predicted by the stimulus physical location proximity, and this resemblance persisted after training. For the delay period, the representational pattern was training dependent.

**Figure 4.**
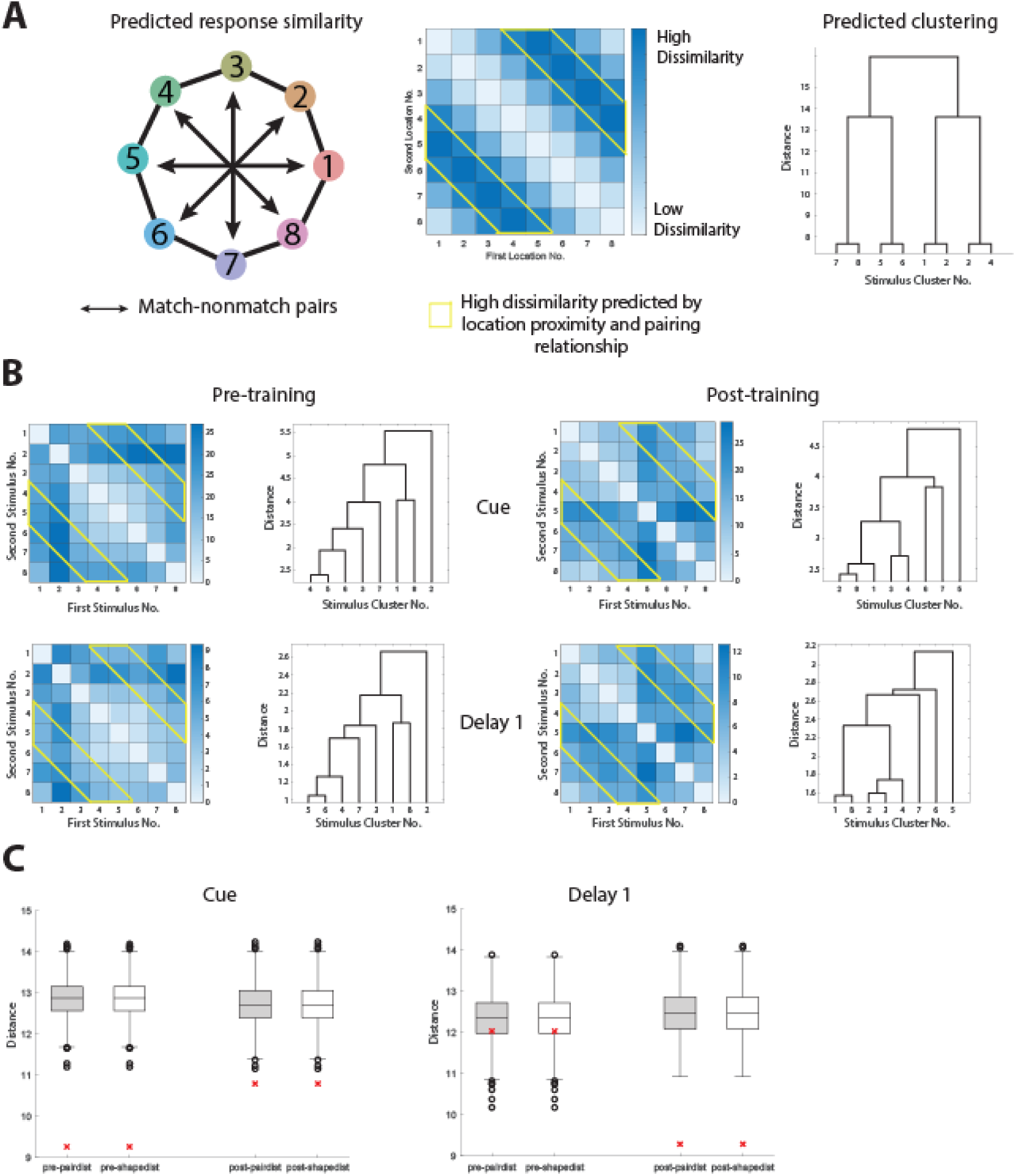
Representational similarity between different spatial locations. A. Diagram, showing the predicted similarity between spatial locations, based on both the spatial proximity and pairing relationship. Heat maps and dendrogram representing the representational distance between any two stimuli used in the spatial task. Stimulus numbers refer to spatial locations as indicated in the diagram. Deeper color in the heat maps indicates less similarity between the neuronal responses elicited by two stimuli. Distance numbers are in arbitrary units. B. measured response similarity between locations in different task epochs. C. Quantification of similarity between the empirical response and the predicted response. The red asterisks are the distances between the dissimilarity matrices in panel B and panel A. Box plots show the distribution of distances after shuffling the stimuli labels for each neuron (empirical vs control, nonparametric test, pre-training.

### Organization of Feature Selectivity

These results of the spatial analysis mostly replicated earlier studies which suggested a local organization of spatial information across the surface of the prefrontal cortex, using different spatial working memory tasks (Constantinidis *et al*. 2001; Leavitt *et al*. 2017). Since it has been argued that spatial representations are fundamentally different than object working memory (Curtis and D’Esposito 2003; Tong and Pratte 2012; D’Esposito and Postle 2015), we were interested to test whether neuronal selectivity for shapes exhibits some organization or not.

We thus repeated our analysis of neuronal selectivity for shape information. The percentage of neurons with significant selectivity for the eight geometric shapes of our dataset evaluated with a 1-way ANOVA test (p<0.05) is shown in Fig. 5. Although shape selectivity was lower compared to spatial selectivity, neurons selective for shape were found both before and after training, with approximately 10% of all neurons recorded exhibiting shape selectivity at any task epoch. We did not find training-dependent selective cell proportion change (Fig.5A), or a change in the degree of selectivity (Fig. 5B; pre-v. post z-score proportion test: cue p=0.063, delay1 p=0.255, sample p=0.560, delay2 p=0.584).

**Figure 5.**
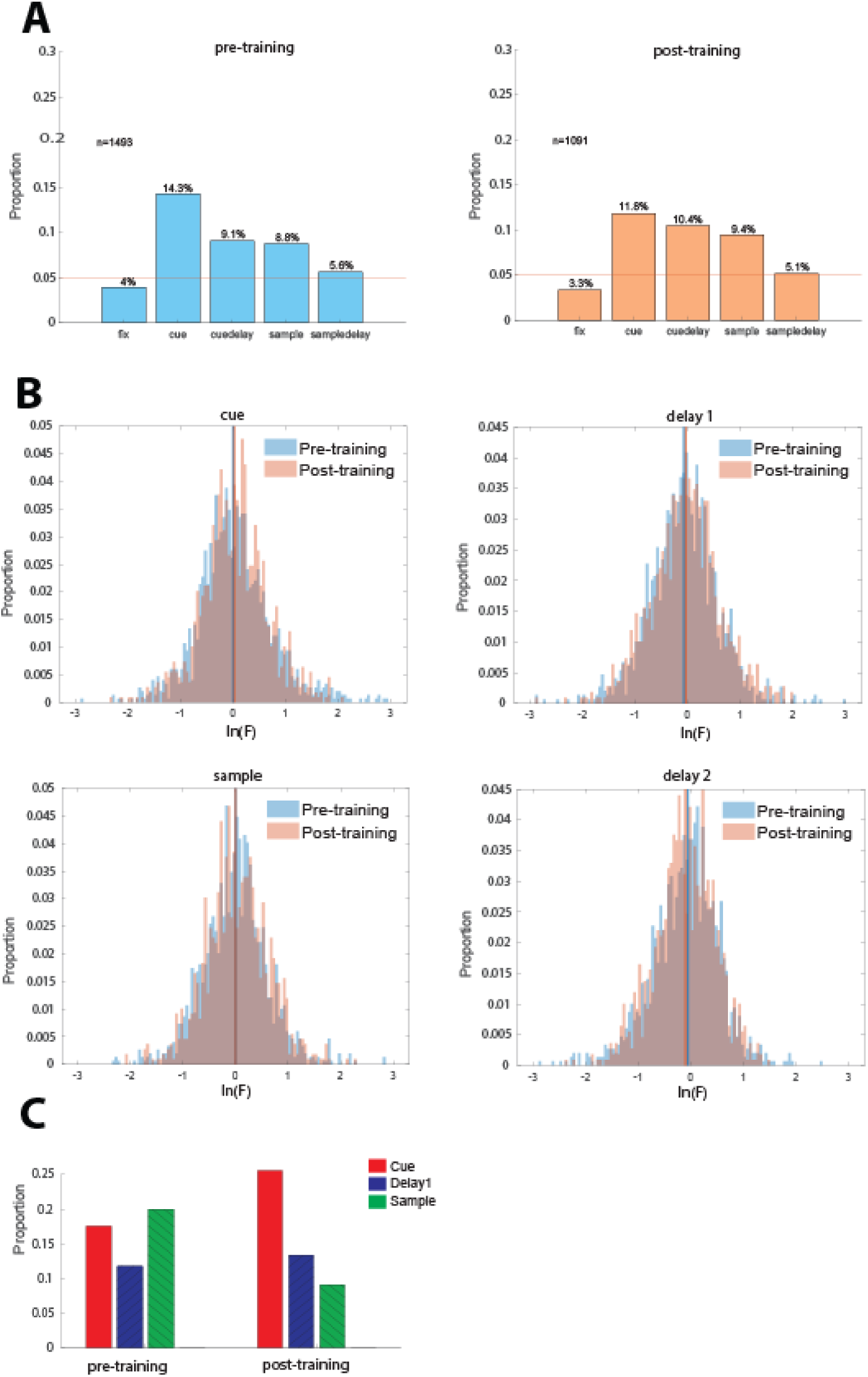
A. Proportion of neurons exhibiting significant shape selectivity at different task epochs. Left. Selective neurons prior to training. Sample size n=1493. Right. Selective neurons post-training (n=1091 neurons). B. Distributions of natural logarithm of F-values in the ANOVA test for shape selectivity. Colored vertical lines represent median of distributions for corresponding training status. C. Proportion of neurons with the same spatial selectivity at different task intervals.

A total of 242 pairs of neurons were recorded simultaneously from separate electrodes spaced 0.2 – 1.5 mm from each other, both neurons of which exhibited shape selectivity in the same task epoch (Fig.5C). We again examined whether these neurons were more likely to exhibit the same preferred stimulus than would be expected by chance. For this dataset, a higher than chance same-shape preference was observed in the cue period, after training (permutation test, p=0.043).

We compared the overall similarity of shape tuning between simultaneously recorded neurons by computing their normalized Euclidean distance, as we did for the spatial stimuli. The mean Euclidean distance was 0.86 prior to training and 0.88 after training. These distances were significantly smaller than would be expected by chance (Fig. 6A; permutation test, p<0.001, in each case). Similar to the spatial task, we also analyzed the spatial clustering of selective cells, regardless of what shape they preferred, and discovered that the chance of finding pairs of selective cells was higher than expected by chance (Fig. 6B-C). Together, these results suggest that shape selectivity too, is not represented randomly across the surface of the cortex. Prefrontal neurons tuned for stimulus shape are present prior to any training as well as after training and organized in patches of cortex with similar shape selectivity.

**Figure 6.**
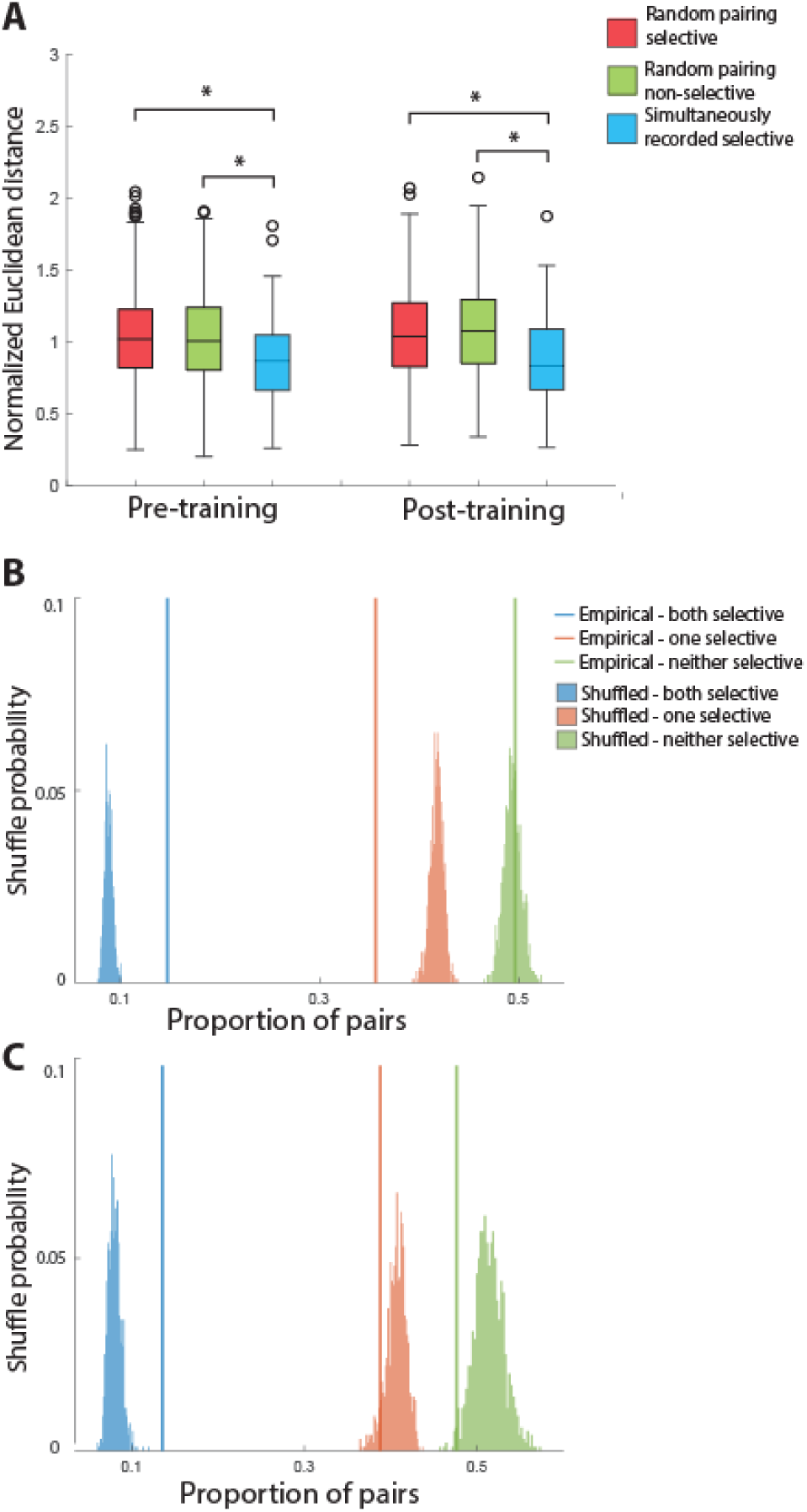
Normalized Euclidean distance of the representations of shape stimuli among pairs of neighboring prefrontal neurons. Conventions are the same as Fig. 3. In the Feature task, the representational distance between nearby neurons is smaller than the expected chance level. B. Permutation test for clustering selective cells in the pre-training fetaure task. Colored vertical lines indicated the proportion of selective/selective, selective/nonselective, nonselective/nonselective pairs drawn from nearby (distance <1.5mm), simultaneously recorded neurons. This empirical proportion was compared to a null distribution constructed by shuffling session labels. C. Selective clustering permutation test for post-training data. Plotting conventions are the same as panel B.

Finally, we determined how different shapes are represented across the population of prefrontal neurons (Fig. 7). Similar to the analysis for the spatial task, we constructed two representational similarity matrices based on the prediction of perceptual similarity of the shapes, as well as based on the prediction of pairing relationship (Fig. 7A, see Methods: Representational similarity). Unexpectedly, after comparing the predictions to the real neural response during various task epochs, we discovered that though the perceptual-based similarity was not able to predict the neural response pattern (shape based prediction: empirical vs control, nonparametric test, pre-training cue p=0.882, post-training cue p=0.990, pre-training delay1 p=0.181, post-training delay1 p=0.676), the pairing-based model performed slightly above chance for the cue period prior to training (Fig. 7C cue period pre-training p=0.014, post-training p=0.685); no significant difference was found for the delay1 period (Fig. 7C pre-training p=0.129, post-training p=0.401).This finding suggests that the statistical structure of sequences of stimuli might influence the organization of the local PFC circuit and this is not tied to task performance, which may itself alter local organization.

**Figure 7.**
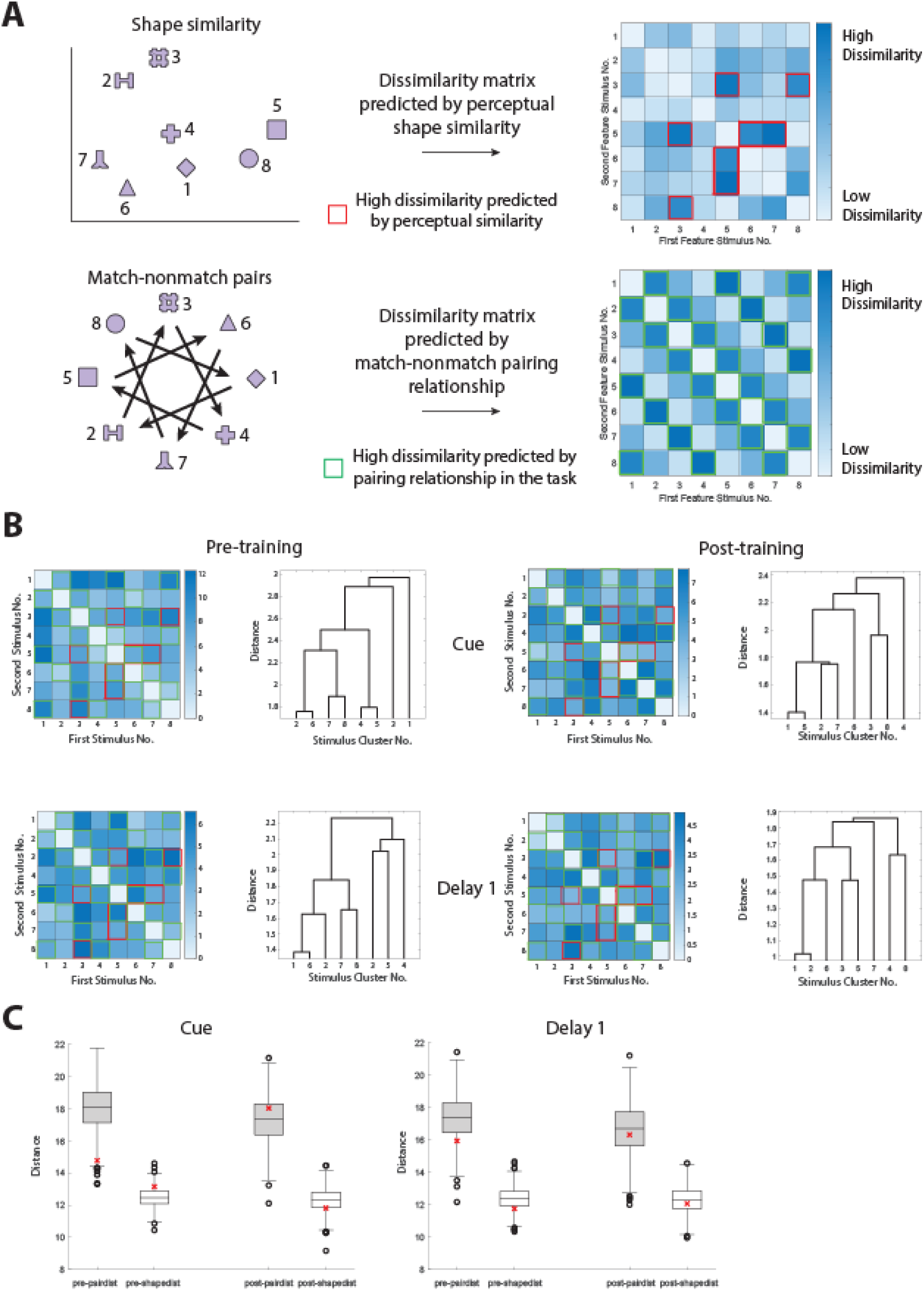
Representational similarity between different shapes. A. Diagram, showing the predicted similarity between spatial locations, based on the spatial proximity (top) and pairing relationship (bottom), respectively. Heat maps and dendrogram representing the representational distance between any two stimuli used in the spatial task. Stimulus numbers refer to spatial locations as indicated in the diagram. Deeper color in the heat maps indicates less similarity between the neuronal responses elicited by two stimuli. Distance numbers are in arbitrary units. B. Measured response similarity between shapes in different task epochs. C. Quantification of similarity between the empirical response and the predicted response. The red asterisks are the distances between the dissimilarity matrices in panel B and panel A (see Methods), The box plots show the distribution of distances after shuffling the stimuli labels for each neuron

## DISCUSSION

Our analysis revealed a local organization of both spatial and non-spatial information across the surface of the prefrontal cortex, with neurons 1.5 mm of each other representing significantly more similar stimuli. These results are consistent with previous studies that have suggested a local organization of spatial representations across the prefrontal cortex, in monkeys trained to perform spatial working memory tasks (Constantinidis *et al*. 2001; Leavitt *et al*. 2017). We now reveal a similar pattern of local organization for object representation, as well. Furthermore, we now document that this organization is largely present even prior to any training to perform a working memory task that required these stimuli to be recognized and maintained in memory.

### Organization of information across the prefrontal cortex

The prefrontal cortex is critical for higher cognitive functions, including the maintenance of information in working memory (D’Esposito and Postle 2015; Riley and Constantinidis 2016). Specifically, this information is thought to be maintained through persistent discharges in a network of brain areas centered on the prefrontal cortex (Constantinidis et al. 2018). Computational models suggest that persistent activity is sustained in neural circuits by virtue of re-entrant connections between neurons with similar tuning for stimulus properties (Compte et al. 2000; Wimmer *et al*. 2014). Neural activity generated by a stimulus presentation is maintained in the network, which behaves as a continuous attractor (Seung et al. 2000). Until now however, the specific organization of these neuronal networks across the surface of the prefrontal cortex has remained unresolved. No obvious, retinotopic organization of visual space has been described across the surface of the prefrontal cortex, or other higher-order cortical areas implicated in the maintenance of working memory, such as the posterior parietal cortex (Qi et al. 2015). Some indications exist for at least some local PFC organization. For example, electrode penetrations that descent into the principal sulcus, parallel to its surface, suggest an orderly progression of receptive field locations (Arnsten 2013). Part of the challenge has been that individual neurons may exhibit non-linear mixed selectivity under different task conditions (i.e. selectivity profiles for the same stimuli that change between experimental conditions) as well as selectivity profiles that differ in different epochs of the same task (Dang et al. 2021). Our current results suggest that selectivity for stimulus properties is an organizing principle in the neural circuits of the prefrontal cortex, as speculated by early models of the cellular basis of working memory (Goldman-Rakic 1995).

Other findings also suggest at least some coarse localization of sensory information across domains. Prefrontal cortex represents information across all sensory modalities and multisensory combinations (Constantinidis 2016; Jaffe and Constantinidis 2021) .Whether object working memory is localized in a different subdivision of the prefrontal cortex than spatial working memory (ventral vs. dorsal) has been a matter of debate (Wilson et al. 1993; Rao *et al*. 1997). In the former viewpoint, dorsal prefrontal cortex is specialized for spatial and ventral prefrontal cortex for object information, which would suggest some type of organization of shape information at least in the ventral prefrontal cortex. In the latter viewpoint, selectivity of individual neurons is predominantly shaped by task demands, with individual neurons in both subdivisions representing types of information required by the task subjects have been trained to perform.

Evaluation of the evidence accrued since these original studies suggests that at least a quantitative difference seems to be evident, with spatial information more prevalent in the dorsolateral prefrontal cortex than the ventrolateral prefrontal cortex (Meyer *et al*. 2011; Constantinidis and Qi 2018). Some organization for the time course of activation has also been revealed across cortical layers: neurons activated by visual stimuli are more likely to be encountered in intermediate layers; neurons with delay period activity in superficial; neurons with motor responses in deep, though this relative segregation is quantitative rather than qualitative (Markowitz et al. 2015; Zhu et al. 2023). Finally, an anterior-posterior gradient of stimulus properties has also been described so that posterior neurons are more selective for the sensory attributes of stimuli no matter what their context, whereas more anterior neurons are much more likely to become activated only in the context of learned tasks (Riley et al. 2017; Riley *et al*. 2018).

Much less is known about the organization principles that might govern representation of object information. Such organization has been discovered in the inferior temporal cortex, with distinct patches specialized for classes of objects, such as faces (Tsao et al. 2006; Dubois et al. 2015; Moeller et al. 2017). Some evidence from imaging studies suggest the existence of functional patches in the prefrontal cortex (Chang et al. 2017). Our current results suggest that shape information in the prefrontal cortex may parallel that of the inferior temporal cortex.

### Training effects

The activity of prefrontal neurons is highly plastic and responding to abstract variables of the task being executed (Fuster et al. 2000; Wallis et al. 2001; Mante et al. 2013). Neurophysiological studies of training have revealed that learning to perform cognitive tasks generally increases responsiveness to stimuli that are relevant for the task execution (Meyers et al. 2012; Constantinidis and Klingberg 2016). Prefrontal neurons, and particularly those in more anterior areas, become responsive to stimuli that previously failed to elicit a response (Qi et al. 2011; Riley *et al*. 2018). Recent human imaging results parallel these findings, confirming appearance of stimulus selectivity for learned stimuli after training (Miller et al. 2022). A variety of other changes have also been reported, including in variability and correlation of prefrontal discharges (Qi and Constantinidis 2012).

In view of these findings, it appeared possible that the local organization of the prefrontal cortex with respect to stimulus selectivity might only emerge after training. Contrary to this expectation, we found neurons with similar selectivity at nearby locations in the cortex, even in naïve monkeys, prior to any training. This result is consistent with previous findings suggesting that underlying network circuitry appears to connect neurons with similar properties, and that these functional modules of neurons remain fairly stable across different behavioral contexts or tasks (Kiani *et al*. 2015).

## ACKNOWLEDGMENTS

Research reported in this paper was supported by the NIH National Eye Institute under award number R01 EY017077 and the NSF under award CRCNS 2011514. We wish to acknowledge Chrissy Suell for technical help; Junda Zhu, and Zhengyang Wang for helpful comments on the manuscript.

## SUPPLEMENTARY FIGURES

**Figure S1.**
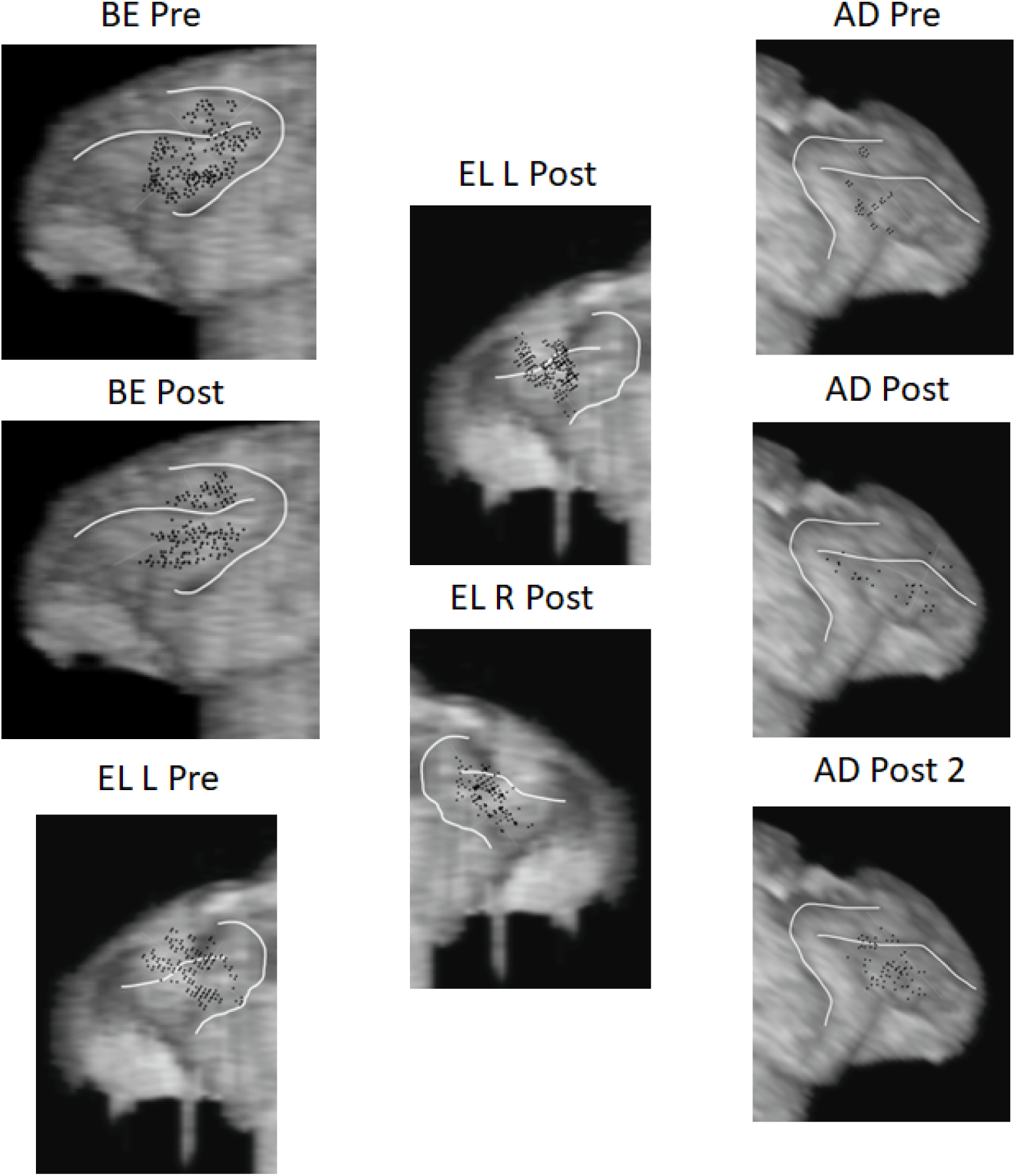
Anatomical localization of electrode penetrations

**Figure S2.**
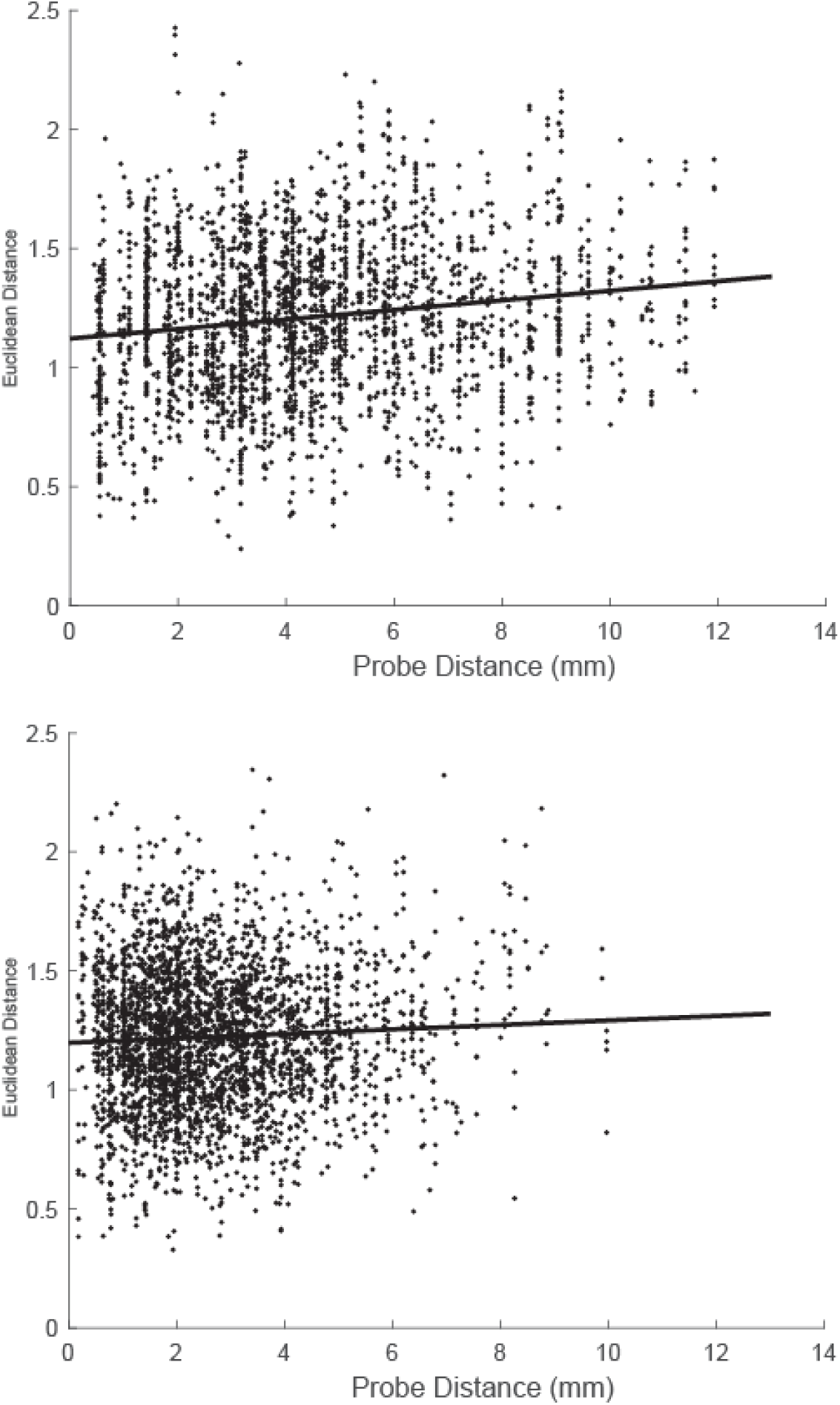
Euclidean distance between the preferred locations of the neurons in every possible pair \as a function distance between the anatomical site where they were recorded (on different days). Top, data from the cue period, prior to training. Black line represents linear regression, p= 6.538e-16. Bottom, data from the cue period after training: p= 3.694e-03.

## REFERENCES

Albers AM, Kok P, Toni I, Dijkerman HC, de Lange FP. 2013. Shared representations for working memory and mental imagery in early visual cortex. Current biology : CB. 23:1427–1431.

Arnsten AF. 2013. The neurobiology of thought: the groundbreaking discoveries of Patricia Goldman-Rakic 1937-2003. Cereb Cortex. 23:2269–2281.

Brainard DH. 1997. The Psychophysics Toolbox. Spat Vis. 10:433–436.

Chang L, Bao P, Tsao DY. 2017. The representation of colored objects in macaque color patches. Nat Commun. 8:2064.

Compte A, Brunel N, Goldman-Rakic PS, Wang XJ. 2000. Synaptic mechanisms and network dynamics underlying spatial working memory in a cortical network model. Cereb Cortex. 10:910–923.

Constantinidis C. 2016. A Code for Cross-Modal Working Memory. Neuron. 89:3–5.

Constantinidis C, Franowicz MN, Goldman-Rakic PS. 2001. Coding specificity in cortical microcircuits: a multiple electrode analysis of primate prefrontal cortex. J Neurosci. 21:3646–3655.

Constantinidis C, Funahashi S, Lee D, Murray JD, Qi XL, Wang M, Arnsten AFT. 2018. Persistent Spiking Activity Underlies Working Memory. J Neurosci. 38:7020–7028.

Constantinidis C, Klingberg T. 2016. The neuroscience of working memory capacity and training. Nat Rev Neurosci. 17:438–449.

Constantinidis C, Qi XL. 2018. Representation of Spatial and Feature Information in the Monkey Dorsal and Ventral Prefrontal Cortex. Front Integr Neurosci. 12:31.

Curtis CE, D’Esposito M. 2003. Persistent activity in the prefrontal cortex during working memory. Trends in Cognitive Sciences. 7:415–423.

D’Esposito M, Postle BR. 2015. The cognitive neuroscience of working memory. Annu Rev Psychol. 66:115–142.

Dang W, Jaffe RJ, Qi XL, Constantinidis C. 2021. Emergence of Nonlinear Mixed Selectivity in Prefrontal Cortex after Training. J Neurosci. 41:7420–7434.

Dubois J, de Berker AO, Tsao DY. 2015. Single-unit recordings in the macaque face patch system reveal limitations of fMRI MVPA. J Neurosci. 35:2791–2802.

Ester EF, Anderson DE, Serences JT, Awh E. 2013. A Neural Measure of Precision in Visual Working Memory. Journal of Cognitive Neuroscience. 25:754–761.

Funahashi S, Bruce CJ, Goldman-Rakic PS. 1989. Mnemonic coding of visual space in the monkey’s dorsolateral prefrontal cortex. J Neurophysiol. 61:331–349.

Fuster JM, Alexander GE. 1971. Neuron activity related to short-term memory. Science. 173:652–654.

Fuster JM, Bodner M, Kroger JK. 2000. Cross-modal and cross-temporal association in neurons of frontal cortex. Nature. 405:347–351.

Goldman-Rakic PS. 1988. Topography of cognition: parallel distributed networks in primate association cortex. Annu Rev Neurosci. 11:137–156.

Goldman-Rakic PS. 1995. Cellular basis of working memory. Neuron. 14:477–485.

Harris KD, Henze DA, Csicsvari J, Hirase H, Buzsaki G. 2000. Accuracy of tetrode spike separation as determined by simultaneous intracellular and extracellular measurements. J Neurophysiol. 84:401–414.

Harrison SA, Tong F. 2009. Decoding reveals the contents of visual working memory in early visual areas. Nature. 458:632–635.

Hart E, Huk AC. 2020. Recurrent circuit dynamics underlie persistent activity in the macaque frontoparietal network. Elife. 9.

Jaffe RJ, Constantinidis C. 2021. Working Memory: From Neural Activity to the Sentient Mind. Compr Physiol. 11:1–41.

Kiani R, Cueva CJ, Reppas JB, Peixoto D, Ryu SI, Newsome WT. 2015. Natural grouping of neural responses reveals spatially segregated clusters in prearcuate cortex. Neuron. 85:1359–1373.

Leavitt ML, Pieper F, Sachs AJ, Martinez-Trujillo JC. 2017. A Quadrantic Bias in Prefrontal Representation of Visual-Mnemonic Space. Cereb Cortex.1–17.

Mante V, Sussillo D, Shenoy KV, Newsome WT. 2013. Context-dependent computation by recurrent dynamics in prefrontal cortex. Nature. 503:78–84.

Markowitz DA, Curtis CE, Pesaran B. 2015. Multiple component networks support working memory in prefrontal cortex. Proc Natl Acad Sci U S A. 112:11084–11089.

Mendoza-Halliday D, Torres S, Martinez-Trujillo JC. 2014. Sharp emergence of feature-selective sustained activity along the dorsal visual pathway. Nat Neurosci. 17:1255–1262.

Meyer T, Constantinidis C. 2005. A software solution for the control of visual behavioral experimentation. J Neurosci Methods. 142:27–34.

Meyer T, Qi XL, Stanford TR, Constantinidis C. 2011. Stimulus selectivity in dorsal and ventral prefrontal cortex after training in working memory tasks. J Neurosci. 31:6266–6276.

Meyers EM, Qi XL, Constantinidis C. 2012. Incorporation of new information into prefrontal cortical activity after learning working memory tasks. Proc Natl Acad Sci U S A. 109:4651–4656.

Miller EK, Erickson CA, Desimone R. 1996. Neural mechanisms of visual working memory in prefrontal cortex of the macaque. J Neurosci. 16:5154–5167.

Miller JA, Tambini A, Kiyonaga A, D’Esposito M. 2022. Long-term learning transforms prefrontal cortex representations during working memory. Neuron. 110:3805–3819 e3806.

Moeller S, Crapse T, Chang L, Tsao DY. 2017. The effect of face patch microstimulation on perception of faces and objects. Nat Neurosci. 20:743–752.

Morgenstern Y, Hartmann F, Schmidt F, Tiedemann H, Prokott E, Maiello G, Fleming RW. 2021. An image-computable model of human visual shape similarity. PLoS Comput Biol. 17:e1008981.

Qi XL, Constantinidis C. 2012. Correlated discharges in the primate prefrontal cortex before and after working memory training Eur J Neurosci. 36:3538–3548.

Qi XL, Elworthy AC, Lambert BC, Constantinidis C. 2015. Representation of remembered stimuli and task information in the monkey dorsolateral prefrontal and posterior parietal cortex. J Neurophysiol. 113:44–57.

Qi XL, Meyer T, Stanford TR, Constantinidis C. 2011. Changes in Prefrontal Neuronal Activity after Learning to Perform a Spatial Working Memory Task. Cereb Cortex. 21:2722–2732.

Rao SC, Rainer G, Miller EK. 1997. Integration of what and where in the primate prefrontal cortex. Science. 276:821–824.

Riley MR, Constantinidis C. 2016. Role of prefrontal persistent activity in working memory. Front Syst Neurosci. 9:181.

Riley MR, Qi XL, Constantinidis C. 2017. Functional specialization of areas along the anterior-posterior axis of the primate prefrontal cortex. Cereb Cortex. 27:3683–3697.

Riley MR, Qi XL, Zhou X, Constantinidis C. 2018. Anterior-posterior gradient of plasticity in primate prefrontal cortex. Nat Commun. 9:3790.

Seung HS, Lee DD, Reis BY, Tank DW. 2000. Stability of the memory of eye position in a recurrent network of conductance-based model neurons. Neuron. 26:259–271.

Sreenivasan KK, Vytlacil J, D’Esposito M. 2014. Distributed and Dynamic Storage of Working Memory Stimulus Information in Extrastriate Cortex. Journal of Cognitive Neuroscience. 26:1141–1153.

Tong F, Pratte MS. 2012. Decoding patterns of human brain activity. Annual review of psychology. 63:483–509.

Tsao DY, Freiwald WA, Tootell RB, Livingstone MS. 2006. A cortical region consisting entirely of face-selective cells. Science. 311:670–674.

Wallis JD, Anderson KC, Miller EK. 2001. Single neurons in prefrontal cortex encode abstract rules. Nature. 411:953–956.

Wilson FA, O Scalaidhe SP, Goldman-Rakic PS. 1993. Dissociation of object and spatial processing domains in primate prefrontal cortex. Science. 260:1955–1958.

Wimmer K, Nykamp DQ, Constantinidis C, Compte A. 2014. Bump attractor dynamics in prefrontal cortex explains behavioral precision in spatial working memory. Nat Neurosci. 17:431–439.

Xing Y, Ledgeway T, McGraw PV, Schluppeck D. 2013. Decoding working memory of stimulus contrast in early visual cortex. The Journal of neuroscience : the official journal of the Society for Neuroscience. 33:10301–10311.

Zhu J, Hammond BM, Constantinidis C. 2023. Laminar pattern of adolescent development changes in working memory neuronal activity. bioRxiv. doi10.1101/2023.07.28.550982.

